# Proteomics support the threespine stickleback egg coat as a protective oocyte envelope

**DOI:** 10.1101/2020.03.04.976316

**Authors:** Emily E. Killingbeck, Damien B. Wilburn, Gennifer E. Merrihew, Michael J. MacCoss, Willie J. Swanson

## Abstract

After the end of the last ice age, ancestrally marine threespine stickleback fish (*Gasterosteus aculeatus*) have undergone an adaptive radiation into freshwater environments throughout the Northern Hemisphere, creating an excellent model system for studying molecular adaptation and speciation. Stickleback populations are reproductively isolated to varying degrees, despite the fact that they can be crossed in the lab to produce viable offspring. Ecological and behavioral factors have been suggested to underlie incipient stickleback speciation. However, reproductive proteins represent a previously unexplored driver of speciation. As mediators of gamete recognition during fertilization, reproductive proteins both create and maintain species boundaries. Gamete recognition proteins are also frequently found to be rapidly evolving, and their divergence may culminate in reproductive isolation and ultimately speciation. As an initial investigation into the contribution of reproductive proteins to stickleback reproductive isolation, we characterized the egg coat proteome of threespine stickleback eggs. In agreement with other teleosts, we find that stickleback egg coats are comprised of homologs to the zona pellucida (ZP) proteins ZP1 and ZP3. We explore aspects of stickleback ZP protein biology, including glycosylation, disulfide bonding, and sites of synthesis, and find many substantial differences compared to their mammalian homologs. Furthermore, molecular evolutionary analyses indicate that *ZP3*, but not *ZP1*, has experienced positive Darwinian selection across teleost fish. Taken together, these changes to stickleback ZP protein architecture suggest that the egg coats of stickleback fish, and perhaps fish more generally, have evolved to fulfill a more protective functional role than their mammalian counterparts. Data are available via ProteomeXchange with identifiers PXD017488 and PXD017489.

## Introduction

Threespine stickleback fish have been called “Darwin’s fishes” in light of their remarkable adaptive radiation throughout the Northern Hemisphere following glacial retreat at the end of the last ice age (∼12,000 years ago) (1, 2). Ancestrally marine fish have colonized thousands of freshwater lakes and streams, evolving significant diversity in morphology, behavior, physiology, and life history (1–4). These divergent forms come into contact with each other, but are frequently reproductively isolated, making stickleback an ideal model system for speciation research (2, 4). Speciation, in the sense of sympatric populations of stickleback coexisting without interbreeding, is often rapid, and attributed to differences in male morphology and behavior and female preferences for those traits as well as ecological selection against hybrids (2, 4, 5). Despite nearly complete reproductive isolation in the wild, however, virtually any stickleback can be crossed in the lab to produce viable, fertile offspring (1). Whereas the evolution of reproductive isolation in stickleback has been attributed to divergent natural and sexual selection, the contribution of rapidly evolving reproductive proteins to stickleback speciation has so far not been considered (4). To begin to address this question from the perspective of female reproductive protein evolution, we have characterized the egg coat proteome of threespine stickleback fish.

Animal oocytes are surrounded by a specialized glycoprotein extracellular matrix termed the “egg coat” (6–8). The egg coat is an interface between the egg and its environment, protecting the oocyte from physical, chemical, and biological hazards (6, 7, 9, 10). It is also an interface between gametes during fertilization, playing roles in attracting and activating sperm, mediating sperm recognition and binding, and blocking the detrimental fitness costs of polyspermy (6, 7, 10). The egg coat goes by different names in the major vertebrate lineages, including the zona pellucida (ZP) in mammals, the vitelline envelope in non-mammals, and the chorion in fish (6, 11). Despite historically complicated nomenclature, egg coats are generally comprised of a common set of glycoproteins characterized by the zona pellucida (ZP) module (12). The ZP module is an ∼260 residue polymerization element consisting of a N-terminal ZP-N domain and a C-terminal ZP-C domain that both adopt immunoglobulin (Ig)-like folds (12–14). Beyond the core ZP module, many ZP proteins have more elaborate structures, including trefoil domains, transmembrane domains, consensus furin protease cleavage sites (CFCS), and tandem arrays of ZP-N repeats that have evolved independently of one another and their associated ZP-C (15–20). Since ZP-N and ZP-C are independent structural domains, we will use the term “ZP module” to refer to the combined ZP-N and ZP-C domains rather than the more generic “ZP domain” (19, 21).

Vertebrate ZP proteins arose from a common ancestral gene through multiple duplication events hundreds of millions of years ago, giving rise to five gene families: *ZP1*/*ZP4*, *ZP2*, *ZP3*, *ZPAX*, and *ZPD* (22, 23). ZP3 proteins, which are typically the smallest ZP protein, contain only the ZP module; this minimal architecture as well as molecular phylogenetics suggest that ZP3 may be most similar to the ancestral ZP protein (6, 11, 24, 25). ZP3 proteins can also have repetitive P/Q residues in relatively short stretches (26). With the exception of ZPD, all other ZP protein families (ZP1/4, ZP2, and ZPAX) contain additional ZP-N domain repeats N-terminal to their ZP module (11, 18). These N-terminal ZP-N domains tend to be less conserved among orthologous proteins of different species (18). ZP1-like proteins typically possess a N-terminal ZP-N domain repeat followed by a P/Q-rich region, a trefoil domain, and a ZP module (18, 26). ZP2 proteins are characterized by multiple N-terminal ZP-N domain repeats prior to their ZP module, and the ZP2 homolog ZPAX has an analogous N-terminal ZP-N domain repeat architecture (18).

ZP proteins are synthesized as precursor polypeptides with a signal sequence (SS) at the N-terminus and a C-terminal propeptide containing a transmembrane domain (TMD) (15, 27, 28). In some fish, however, the TMD is absent (26, 28). The ZP module itself consists of 8, 10, or 12 disulfide-bonded cysteine residues, followed by a CFCS and, if present, a TMD or hydrophobic sequence (26–28). The dimerization of ZP-N domains between ZP modules facilitates the assembly of the filamentous egg coat ultrastructure (20, 29–32).

In mammalian egg coats, ZP proteins serve as both structural and sperm-binding proteins (14, 33–38). In fish, however, the role of ZP proteins in the egg coat is less well characterized and may be purely structural (26, 28, 39). Teleost fish sperm lack an acrosome, a secretory vesicle involved in sperm-egg binding, and teleost fish eggs contain an additional structure called the micropyle, a funnel-shaped, narrow channel through the egg coat that permits sperm to reach the plasma membrane of the egg (7, 40–43). The micropyle attracts sperm by chemotaxis, and its precise diameter restricts polyspermy by allowing passage of only one sperm at a time (41, 42, 44, 45). Whereas sperm in other animals bind to and dissolve the egg coat at the point of contact, in teleost fish the micropyle is solely responsible for sperm entry through the egg coat (42).

In mammals, ZP proteins are synthesized in the ovary by oocytes and/or their surrounding follicle cells (25). In fish, however, ZP proteins can be expressed in the liver as well as the ovary in response to estrogen, and subsequently transported through the bloodstream to the ovary to assemble around eggs (9, 46–48). This additional site of ZP synthesis may reflect the comparatively large size of fish egg clutches, necessitating the synthesis of large amounts of protein in a relatively short time (9, 26, 49, 50). ZP1 and ZP3, the most common ZP proteins in fish egg coats, both have paralogous classes of genes with hepatic and ovarian expression (23, 26, 49). Species-specific gene amplifications and losses have resulted in some teleost fish, such as zebrafish, retaining only ovarian expression; others retain both ovary and liver expression, and others solely liver (47, 48). One of the two expression sites typically becomes dominant, with liver synthesis of ZP proteins most common across teleosts (47, 48). Vitellogenin, an egg yolk precursor protein, shows similar hepatic expression and migration to the ovary in the bloodstream of fish, amphibians, and birds (28, 46). In fish, ZP synthesis and vitellogenesis occur simultaneously in response to 17β-estradiol production by follicle cells (9, 28, 39, 47).

Reproductive proteins that mediate gamete recognition during fertilization show species-specificity in both their structure and binding affinities (51, 52). Despite their central role in fertilization, however, reproductive proteins are frequently among the most rapidly evolving genes in any taxa (51, 53–58). This juxtaposition of rapid evolution and functional constraint suggests a role for positive Darwinian selection in the maintenance of sperm-egg interactions. Furthermore, the molecular evolutionary history of a protein can identify sites under adaptive evolution that may be functionally important (59–65). Signatures of rapid, adaptive evolution characteristic of reproductive proteins suggest that sequence diversification can be beneficial to genes involved in reproduction (51). More formally, when nonsynonymous (*d_N_*) substitutions outweigh synonymous (*d_S_*) substitutions, *d_N_*/*d_S_* (also denoted ω) is greater than one and suggests there was positive selection for changes in amino acid sequence (51, 59, 66). Positive selection on gamete recognition proteins can contribute to reproductive isolation between diverging taxa, with variation between diverging populations creating species barriers that may ultimately lead to speciation (6, 7, 54, 55, 67).

Stickleback fish have been very successful vertebrate models of adaptive evolution and speciation, but reproductive proteins have so far not been explored in this teleost speciation system (5, 68). Since reproductive proteins represent some of the best-known examples of adaptive evolution and speciation at the molecular level, we combine proteomics and molecular evolutionary analyses to begin to address this open question (65).

## Experimental Procedures

### Animal statement

Threespine stickleback fish were collected from a single freshwater site in Lake Union, Washington, USA (47°38’55” N, 122°20’47” W) during their annual breeding season in 2015 (Washington Department of Fish and Wildlife permit 15-033 to C. Peichel). Fish were collected with minnow traps, eggs were obtained from gravid females, and they were euthanized shortly after collection by immersion in 0.2% MS-222. All animals were collected under permits obtained from the Washington Department of Fish and Wildlife, and all animal methods were conducted in accordance with the guidelines of the Fred Hutchinson Cancer Research Center Institutional Animal Care and Use Committee (Protocol 1575 to C. Peichel).

### Egg coat isolation

Eggs were obtained from gravid female stickleback by gentle abdominal squeezing, and lysed by periodic homogenization in 1% Triton X-100 detergent in Hank’s solution (138 mM sodium chloride, 5 mM potassium chloride, 0.25 mM sodium phosphate dibasic, 0.4 mM potassium phosphate monobasic, 1.3 mM calcium chloride, 1 mM magnesium sulfate, 4 mM sodium bicarbonate; adapted from (69)). Insoluble egg coats were isolated by centrifugation (2,000 *x g* for 10 minutes), and contaminating egg cytosolic proteins were removed with repeated washes of 1% Triton X-100 in Hank’s solution followed by centrifugation. Some samples were additionally treated with 7 M urea to remove trace contaminants of vitellogenin without affecting major egg coat proteins (Figures 1 and 2).

**Figure 1:**
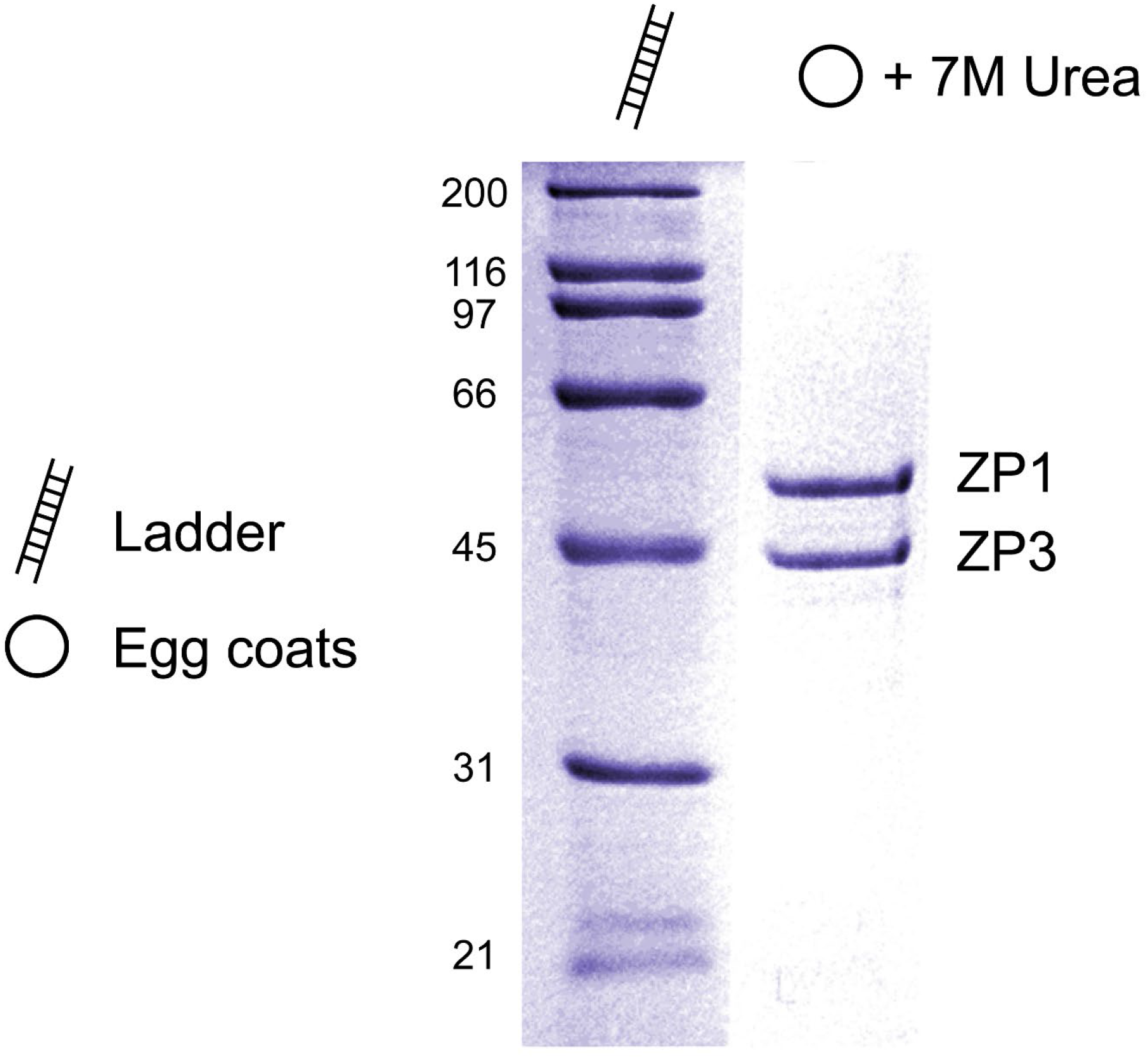
SDS-PAGE of stickleback egg coats. Stickleback egg coats were treated with 7 M urea to remove contaminating vitellogenin, likely carried over from egg coat isolation, and separated by SDS-PAGE. Individual bands were excised from the gel and analyzed by LC-MS/MS, with ZP1 and ZP3 identified as the two major protein components of stickleback egg coats.

**Figure 2:**
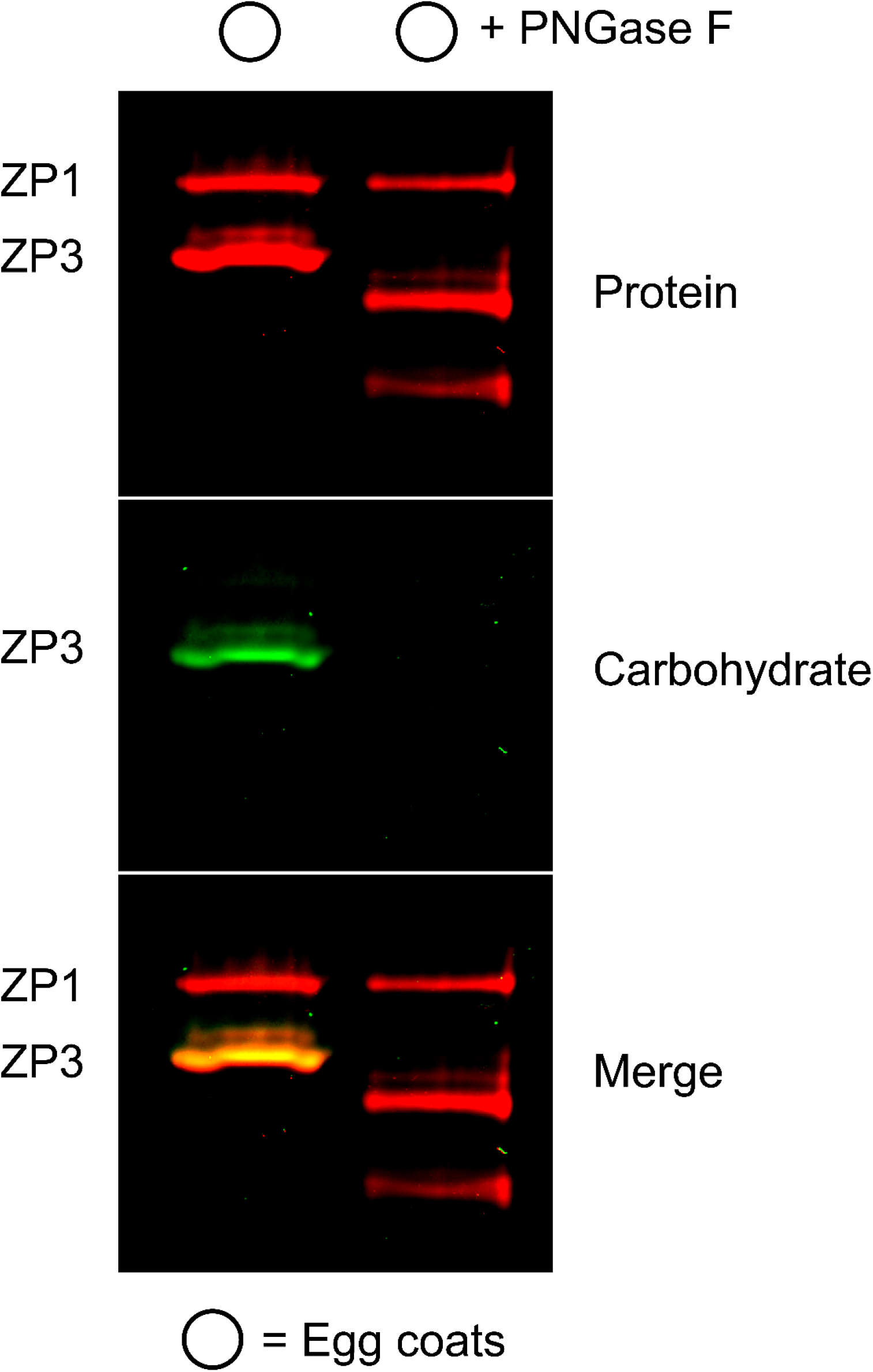
Glycosylation analysis of stickleback egg coats. Stickleback egg coats were treated with 7 M urea, deglycosylated with PNGase F, and separated by SDS-PAGE. The gel was stained with fluorescent carbohydrate and protein dyes, and the images were overlaid to visualize glycoprotein staining. Of the two stickleback egg coat proteins, only ZP3 is glycosylated, and the glycosylation appears to be *N*-linked as the protein no longer stains for carbohydrate after PNGase F treatment. Based on the mass shift after deglycosylation and the single *N*-linked glycosylation site in ZP3, the glycan appears to be ∼4 kDa. Note that the low molecular weight band in the egg coat + PNGase F lane is PNGase F.

### Analysis of egg coats by SDS-PAGE

Stickleback egg coats were analyzed under both reducing and non-reducing conditions by SDS-PAGE with 12% acrylamide gels and a tris-tricine buffering system; electrophoresis was performed at 50 V for 15 minutes, followed by 100 V for 90 minutes (70). Samples were prepared by incubation of solid egg coats in a 1% SDS solution, with or without 2-mercaptoethanol, at 95°C, with insoluble material removed by centrifugation. Proteins were stained with either Coomassie Brilliant Blue R-250 (MilliporeSigma, St. Louis, MO, USA) or SYPRO Ruby (Thermo Fisher Scientific, Waltham, MA, USA). Glycosylation was detected by in-gel periodic acid-Schiff staining using the Pro-Q Emerald 488 glycoprotein staining kit (Invitrogen, Carlsbad, CA, USA) and imaged using a Typhoon FLA 9000 laser bed scanner (GE Healthcare Bio-Sciences, Pittsburgh, PA, USA). To determine if the observed glycosylation of stickleback ZP3 was *N*-linked, egg coats were treated with PNGase F (New England BioLabs, Ipswich, MA, USA) prior to electrophoresis following the manufacturer’s protocol.

### Mass spectral characterization of egg coats

Following SDS-PAGE of stickleback egg coats, individual protein bands were excised using a sterile scalpel blade, cut into ∼1 mm^3^ cubes, and placed in a 1.7 ml tube. Remaining Coomassie dye was extracted through multiple rounds of addition of 50 mM ammonium bicarbonate (with 15 minute incubation), addition of acetonitrile (with 15 minute incubation), removal of supernatant, and drying of the gel pieces in a vacuum centrifuge. After the dye was completely removed, disulfide bonds were reduced by incubating the gel pieces in 20 mM DTT in 100 mM ammonium bicarbonate at 56°C for 45 minutes, followed by alkylation with 55 mM iodoacetamide in 100 mM ammonium bicarbonate in the dark for 30 minutes. The gel pieces were washed twice with 100 mM ammonium bicarbonate, dehydrated with acetonitrile, and incubated with 1 μg trypsin (Promega, Madison, WI, USA) in 50 mM ammonium bicarbonate overnight at 37°C. The supernatant was then collected, the gel pieces washed twice with 50 mM ammonium bicarbonate and acetonitrile, and the washes added to the collected supernatant. The final collected solution was concentrated by evaporative centrifugation and resolubilized in 10 μl 5% acetonitrile / 0.1% formic acid. Three µl of each sample was loaded onto a 30 cm fused silica 75 µm column and 3.5 cm 150 µm fused silica KASIL 1 frit trap (PQ Corporation, Malvern, PA, USA) loaded with 4 µm Jupiter C12 Proteo reverse-phase resin (Phenomenex, Torrance, CA, USA) and analyzed with Thermo Fisher Scientific EASY-nLC. Buffer A was 0.1% formic acid in water and Buffer B was 0.1% formic acid in acetonitrile. The 60-minute LC gradient consisted of 2 to 40% B in 30 minutes, 40 to 60% B in 10 minutes, 60 to 95% B in 5 minutes, followed by a 15 minute wash and a 15 minute column equilibration. Peptides were eluted from the column and electrosprayed into a Velos Pro Linear Ion Trap Mass Spectrometer (Thermo Fisher Scientific). Data was acquired using data-dependent acquisition (DDA) and analyzed using an in-house version of COMET (71, 72) (with a differential modification of 15.994915 Da for methionine and a static modification of 57.021461 Da for cysteine) for database searching against publicly available stickleback ESTs (retrieved from the UCSC Genome Browser) that were assembled with Trinity (73, 74) and six-frame translated. The search database also contained known contaminants such as trypsin and human keratin. Percolator v.2.09 (75) was used to filter the peptide-spectrum matches with a q-value threshold of ≤ 0.01, and peptides were assembled into protein identifications using an in-house implementation of IDPicker (76).

### Sequencing of stickleback ZP cDNA

Total RNA was isolated from *G. aculeatus* ovary and liver tissue by lysis in guanidinium isothiocyanate and cesium chloride gradient ultracentrifugation (procedure modified from (77)). Briefly, tissues were homogenized in five volumes of 4 M guanidinium isothiocyanate in a Dounce homogenizer, 10% SDS was added to a final concentration of 0.1%, and the mixture centrifuged for 5 minutes at 5,000 *x g* to remove insoluble debris. The supernatant was then layered over 5.7 M cesium chloride, centrifuged at 154,000 *x g* for 23 hours at 20°C, purified RNA was washed three times with 70% ethanol, and resuspended in RNase-free water. *G. aculeatus* ovary and liver cDNA was prepared from total RNA using the SMARTer cDNA synthesis kit (Clontech, Palo Alto, CA, USA).

ZP1 and ZP3 coding sequences were PCR amplified from *G. aculeatus* liver cDNA (primer sequences in Table S1), cloned into the pCR4-TOPO vector (Invitrogen), transformed into NEB 5-alpha chemically competent *Escherichia coli* (New England BioLabs), and submitted for Sanger sequencing (Eurofins Genomics, Louisville, KY, USA). Sequences were analyzed using the Lasergene DNASTAR package (v.11.1.0; Madison, WI, USA).

### Disulfide bond characterization

To investigate the disulfide bonding pattern of stickleback ZP1 and ZP3, egg coat samples were prepared under different reduction and alkylation conditions prior to trypsin proteolysis and mass spectral characterization: (1) no reduction or alkylation (disulfide identification), (2) alkylation without reduction (reduced cysteine identification), (3) alkylation followed by reduction (disulfide identification with potentially better trypsin cleavage site accessibility), and (4) reduction followed by alkylation (traditional peptide fingerprinting). To volumetrically match samples, 100 mM ammonium bicarbonate was substituted in place of reagents as necessary. Briefly, an initial reduction was performed with 100 mM BME in 7 M urea in 100 mM ammonium bicarbonate, and the samples were incubated at 60°C for 45 minutes. Samples were then alkylated with 200 mM iodoacetamide in 7 M urea in 100 mM ammonium bicarbonate and incubated for 45 minutes in the dark. A final reduction was performed with 400 mM BME, and all four samples were diluted 1:4 with ammonium bicarbonate to reduce urea concentration. Trypsin (2 µg; Promega) was added to the samples before incubation at 37°C overnight. The samples were then acidified with 1% TFA, desalted by C18 ZipTip (MilliporeSigma), concentrated by evaporative centrifugation, and resolubilized in 10 μl 5% acetonitrile / 0.1% formic acid. Three µl of each sample was loaded onto a 30 cm fused silica 75 µm column and 3.5 cm 150 µm fused silica KASIL 1 frit trap (PQ Corporation) loaded with 3 µm Reprosil-Pur C18 reverse-phase resin (Dr. Maisch, Ammerbuch, Germany) and analyzed with Thermo Fisher Scientific EASY-nLC. Buffer A was 0.1% formic acid in water and buffer B was 0.1% formic acid in acetonitrile. The 100-minute LC gradient consisted of 0 to 16% B in 15 minutes, 16 to 35% B in 60 minutes, 35 to 75% B in 15 minutes, 75 to 100% B in 5 minutes, followed by a 5 minute wash and a 25 minute column equilibration. Peptides were eluted from the column on a 50°C heated source (CorSolutions, Ithaca, NY, USA) and electrosprayed into an Orbitrap Fusion Lumos Mass Spectrometer (Thermo Fisher Scientific). Data was acquired using data-dependent acquisition (DDA) with dynamic exclusion turned off. Mass spectral data was analyzed with MassMatrix v.3.0.10.25 to detect disulfide-linked peptides, with ZP1 and ZP3 coding sequences (cloning described above) used as the search database (78–83).

### Molecular evolution of teleost ZP proteins

To assess ZP gene evolution within teleost fish, 30 species were chosen spanning the teleost phylogeny, with ZP1 and ZP3 open reading frames (ORFs) identified by homology to stickleback ZP1 and ZP3 using TBLASTX (84, 85). For the 31 total species (including *G. aculeatus*), sequences for each gene were aligned separately using Clustal Omega and a concatenated gene tree was constructed using RAxML with the PROTGAMMALG substitution model (Figure S1 and supplemental materials; (86, 87)). Rates of molecular evolution were calculated using PAML v.4.8, with site models M8a (nearly neutral) and M8 (positive selection) compared by likelihood ratio test (88, 89). Sites under positive selection were defined as coding positions with a Bayes empirical Bayes (BEB) posterior probability of > 50% under M8 (90).

A homology model of stickleback ZP3 was generated using Rosetta by threading of the stickleback ZP3 sequence to the available chicken ZP3 structure (PDB ID: 3NK4; (13)) (aligned using Clustal Omega), loop modeling using cyclic coordinate descent (CCD) with refinement by kinetic closure (KIC), and full atom minimization using the relax function (87, 91, 92). *N*-glycosylation was modeled using GlycanBuilder (93).

### Experimental Design and Statistical Rationale

Initial biochemical investigations of stickleback egg coats found no detectable differences among the females sampled (biological replicates). To minimize the number of breeding females that were trapped and euthanized, we did not use replicates, with the exception of the glycosylation assay where technical replicates (n = 2) were employed. In total, egg coat samples from three females were used for the analyses described in this manuscript.

## Results

### Egg coat glycoprotein characterization

To characterize the proteome of threespine stickleback egg coats, egg coats were isolated and examined by SDS-PAGE (Figure S2). Individual bands were excised and analyzed by liquid chromatography-tandem mass spectrometry (LC-MS/MS), with the two main protein components of stickleback egg coats identified as ZP1 and ZP3 (Table S2). The remaining bands represent carryover of vitellogenin from the egg yolk during egg coat isolation. Treatment with 7 M urea removes the contaminating vitellogenin bands, with no apparent loss in intensity of ZP1 or ZP3 (Figure 1).

Reproductive proteins are frequently glycosylated (94, 95). These post-translational modifications affect protein solubility and stability, and are thought to play a role in gamete recognition (59, 96). Glycosylation analysis of stickleback egg coats indicates that of the two main egg coat proteins, only ZP3 is glycosylated (Figure 2). Stickleback ZP3 has a single putative *N*-glycosylation motif at N_160_, and treatment with PNGase F confirmed that the glycan is *N*-linked. Notably, this particular *N*-linked glycosylation site is highly conserved from fish to mammals (97).

### ZP disulfide bond characterization

The insoluble nature of stickleback egg coats in the absence of reducing conditions (even in 7 M urea) suggests that intermolecular disulfide bonds may stabilize the egg coat structure. To determine the disulfide bonding patterns of stickleback ZP1 and ZP3, egg coats were subjected to differing reduction and alkylation conditions prior to performing LC-MS/MS, with dynamic exclusion turned off to permit more quantitative peptide spectral counting. Reverse Transcriptase PCR (RT-PCR) of ZP1 and ZP3 from stickleback ovary and liver RNA produced bands from liver RNA only, suggesting that in stickleback ZP genes are transcribed in the liver (Figure S3). ZP sequences obtained from RT-PCR were used as the database for LC-MS/MS searches, resulting in 66% sequence coverage from 25 peptides for ZP1 and 69% sequence coverage from 24 peptides for ZP3.

ZP proteins have characteristic disulfide bonding patterns within the ZP-N and ZP-C domains of their ZP modules. The crystal structure of chicken ZP3, for instance, shows a C_1_-C_4_, C_2_-C_3_ connectivity for ZP-N and a C_5_-C_7_, C_6_-C_11_, C_8_-C_9_, C_10_-C_12_ connectivity for ZP-C (PDB ID: 3NK4; (13)). Although our analysis generally found evidence of homologous disulfide bonding in stickleback ZP proteins, we also see evidence for shuffled disulfides and new cysteines that could alter the disulfide bonding of stickleback ZP proteins (summarized in Figure 3). For both ZP1 and ZP3, the majority of cysteines within the ZP module were modifiable with iodoacetamide in the absence of reducing agent, suggesting variable and/or transient disulfide bonding. Consistent with disulfide shuffling, both stickleback ZP1 and ZP3 have an odd number of cysteine residues in their ZP module (11 vs. 12 in chicken ZP3). Furthermore, stickleback ZP1 has two additional cysteines (C_4_ and C_5_) in the linker between its ZP-N and ZP-C domains, and ZP3 has an additional cysteine (C_9_) in its ZP-C domain (see Figure 3). Table 1 provides counts for all potential disulfide-linked peptides by LC-MS/MS for ZP1, and Table 2 provides counts for ZP3.

**Figure 3:**
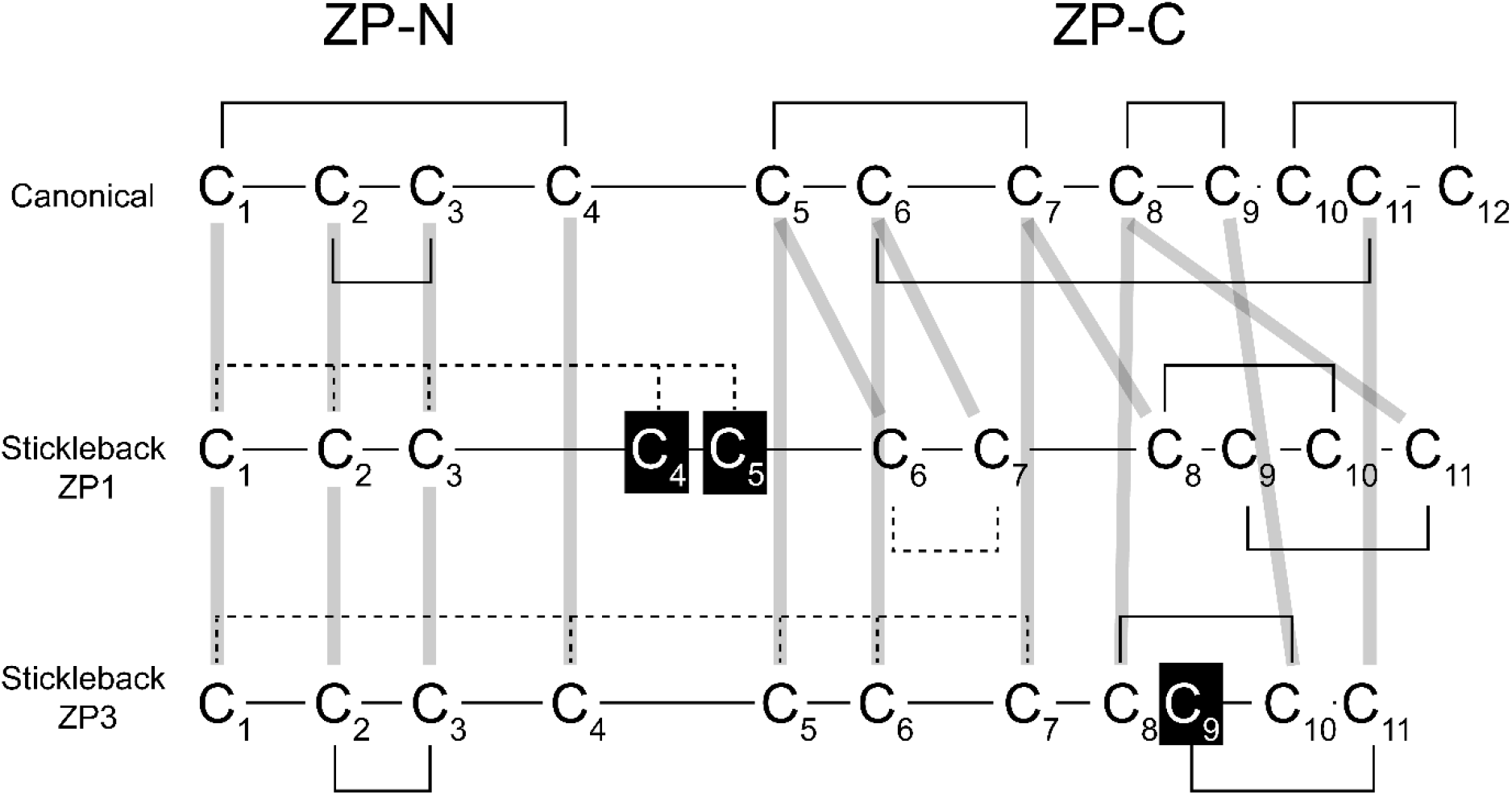
Disulfide bonding pattern of stickleback ZP proteins. Summary of the observed disulfide bonding pattern of stickleback ZP1 and ZP3. The “canonical” disulfide bonding pattern is from the crystal structure of chicken ZP3 (PDB ID: 3NK4; (13)), with cysteines connected by grey lines homologous between stickleback and chicken ZP3. Relative distance between cysteines is indicated by the length of the backbone. Dashed disulfide bonds in stickleback ZP1 and ZP3 denote potential disulfide shuffling, as the indicated cysteines were found to be carbamidomethyl (CAM) modified by mass spectrometry. Cysteines 4 and 5 of stickleback ZP1 (boxed in black) and cysteine 9 of stickleback ZP3 (boxed in black) have no homolog in chicken. Note that cysteines 9 and 10 of stickleback ZP1 are not homologous to chicken ZP3, but do have homologs in chicken ZP1, so are not indicated in black. Stickleback ZP1 is additionally missing its canonical cysteine 4 in its ZP-N domain. For both stickleback ZP1 and ZP3, cysteines 8, 9, 10, and 11 were present on the same peptide so disulfide bonding was inferred by homology to chicken.

**Table 1:** Stickleback ZP1 disulfide bonding patterns by mass spectrometry. The disulfide bonding pattern of stickleback ZP1 was assessed by detection of cysteine cross-linked peptides with LC-MS/MS; the number of peptides supporting each disulfide bond are indicated in parentheses. Homologous cysteines are from a chicken ZP3 crystal structure (PDB ID: 3NK4; (13)), as no ZP1 structure is currently available; a dash indicates that no homologous cysteine is present in chicken ZP3. Cysteines are considered “IAA reactive” if they were found to be carbamidomethyl (CAM) modified after iodoacetamide (IAA) alkylation. Disulfide bonding patterns were inferred by homology to chicken for cysteines 6 and 7, as no cross-linked peptides were detected, as well as for cysteines 8, 9, 10, and 11, since they were present on the same peptide. Note that stickleback ZP1 contains an odd number of cysteine residues.

| Stickleback cysteine number | Homologous cysteine in chicken | IAA reactive | Potential disulfide bonds |
| --- | --- | --- | --- |
| 1 | 1 | ✓ | 2 (5) |
| 2 | 2 | ✓ | 1 (5), 4 (6) |
| 3 | 3 | ✓ | 4 (4) |
| 4 | – | ✓ | 2 (6), 3 (4), 5 (23) |
| 5 | – | ✓ | 4 (23) |
| 6 | 5 | ✓ | 7 (none detected) |
| 7 | 6 | ✓ | 6 (none detected) |
| 8 | 7 |  | 10 (3) |
| 9 | – |  | 11 (3) |
| 10 | – |  | 8 (3) |
| 11 | 8 |  | 9 (3) |

**Table 2:** Stickleback ZP3 disulfide bonding patterns by mass spectrometry. The disulfide bonding pattern of stickleback ZP3 was assessed by detection of cysteine cross-linked peptides with LC-MS/MS; the number of peptides supporting each disulfide bond are indicated in parentheses. Homologous cysteines are from a chicken ZP3 crystal structure (PDB ID: 3NK4; (13)); a dash indicates that no homologous cysteine is present in chicken ZP3. Cysteines are considered “IAA reactive” if they were found to be carbamidomethyl (CAM) modified after iodoacetamide (IAA) alkylation. Cysteines 8, 9, 10, and 11 were present on the same peptide, so bonding patterns of these cysteines were inferred by homology to chicken; additionally, cysteines 8 and 9 are consecutive residues and cannot disulfide bond with each other. Note that stickleback ZP3 contains an odd number of cysteine residues.

| Stickleback cysteine number | Homologous cysteine in chicken | IAA reactive | Potential disulfide bonds |
| --- | --- | --- | --- |
| 1 | 1 | ✓ | 4 (24), 5 (5), 6 (30), 7 (2) |
| 2 | 2 |  | 3 (6) |
| 3 | 3 |  | 2 (6) |
| 4 | 4 | ✓ | 1 (24), 5 (10), 6 (11) |
| 5 | 5 | ✓ | 1 (5), 4 (10), 6 (19) |
| 6 | 6 | ✓ | 1 (30), 4 (11), 5 (19) |
| 7 | 7 | ✓ | 1 (2) |
| 8 | 8 |  | 10 (9) |
| 9 | – |  | 11 (9) |
| 10 | 9 |  | 8 (9) |
| 11 | 11 |  | 9 (9) |

### ZP molecular evolution

Molecular evolutionary analyses of stickleback *ZP1* and *ZP3* suggest that the divergence of *ZP3* across teleosts has been driven by positive Darwinian selection, with 2.3% of sites in *ZP3* under positive selection with ω = 1.89. To test for selection, a model of positive selection (M8) was compared to a model of neutral evolution (M8a) by likelihood ratio test (88, 89). These nested models allow for variation in ω among codons, but the null model M8a restricts ω to 1 while the alternative model M8 allows for adaptive evolution with ω > 1. For *ZP3*, M8 fits the data significantly better than M8a, suggesting that allowing sites with ω > 1 significantly improves the fit of the model to the data (p = 1.2 x 10^-4^; parameters summarized in Table 3). A similar test for adaptive evolution of *ZP1* across teleosts was not significant, in agreement with previous work where *ZP3* has been found to be under selection in mammals while *ZP1* is not. To our knowledge, this was the first investigation of ZP molecular evolution in fish (54, 98, 99). Residues under selection in stickleback ZP3 are indicated as red spheres in Figure 4.

**Figure 4:**
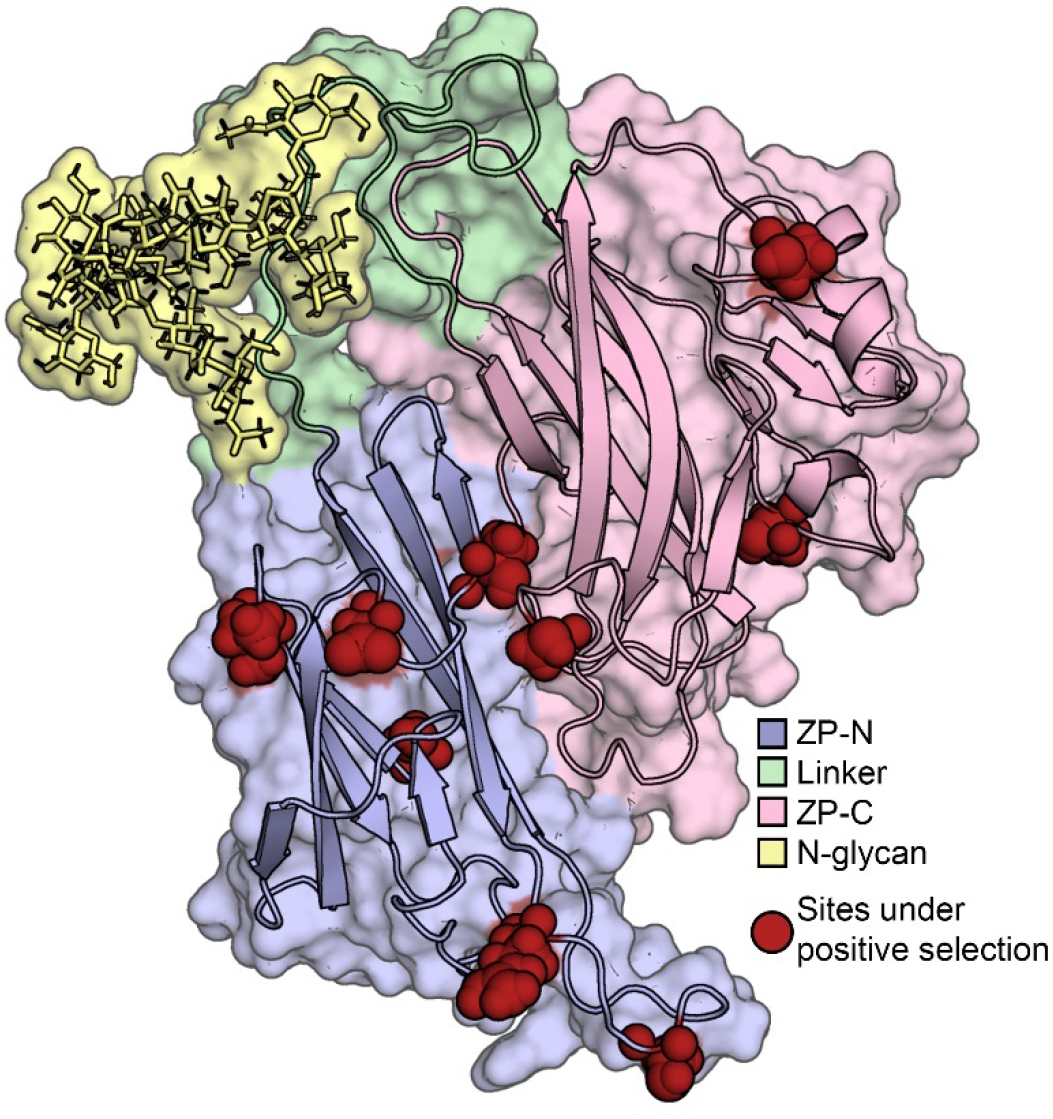
Stickleback ZP3 homology model. Homology model of stickleback ZP3. ZP-N and ZP-C domains are colored purple and pink, respectively; the linker between the domains is indicated in green; residues under positive selection across teleosts are denoted as red spheres; the single *N*-linked glycosylation site in stickleback ZP3 is shown in yellow. Note that sites under positive selection in *ZP3* tend to cluster, particularly those within the ZP-N domain.

**Table 3:** Evolutionary rate analysis of stickleback ZP proteins. The proportion of sites under positive selection (*p*_1_) or under selective constraint (*p*_0_) and the parameters *p* and *q* for the beta distribution *B*(*p*, *q*) are given for *ZP1* and *ZP3* across teleosts. P-values for a likelihood ratio test comparing M8 (selection) to M8a (nearly neutral) are shown, with significant results highlighted in bold (88, 89). Sites under selection in *ZP3* are specified with respect to stickleback, with the signal peptide included.

| Egg coat protein | M8a (neutral model) | M8 (positive selection) | $-2\Delta\log L$ | p-value | Sites under selection |
| --- | --- | --- | --- | --- | --- |
| **ZP1** | $p_0 = 0.88429$,<br>$p = 0.72988$,<br>$q = 2.60577$,<br>$p_1 = 0.11571$,<br>$\omega = 1$ | $p_0 = 0.88920$,<br>$p = 0.72148$,<br>$q = 2.52174$,<br>$p_1 = 0.11080$,<br>$\omega = 1.02158$ | 0.071964 | 0.39 | – |
| **ZP3** | $p_0 = 0.94033$,<br>$p = 0.65445$,<br>$q = 1.85936$,<br>$p_1 = 0.05967$,<br>$\omega = 1$ | $p_0 = 0.97745$,<br>$p = 0.61635$,<br>$q = 1.52432$,<br>$p_1 = 0.02255$,<br>**$\omega = 1.88757$** | 13.5158 | **$1.2 \times 10^{-4}$** | 76, 86, 132,<br>136, 155,<br>159, 256,<br>283, 329 |

## Discussion

As the first barrier sperm encounter during fertilization, the egg coat is an essential determinant of reproductive isolation in any taxa (52, 55, 100). Egg coat proteins are frequently rapidly evolving, and their divergence contributes to reproductive isolation and suggests a role in speciation (6, 7, 51, 52, 54–56, 67, 101, 102). With shotgun proteomics using liquid chromatography-tandem mass spectrometry (LC-MS/MS), we find that the stickleback egg coat is comprised of homologs to the zona pellucida (ZP) glycoproteins ZP1 and ZP3 (Figure 1). Our findings are consistent with egg coat characterization in other fish, where ZP1, ZP3, and occasionally the ZP2 homolog ZPAX are the main structural proteins (26, 49, 97).

Egg coat glycoproteins, as with other reproductive proteins, are frequently glycosylated (94, 95). These carbohydrate modifications are thought to be involved in gamete recognition during fertilization, and to contribute to egg coat solubility (59, 96, 103, 104). Of the two stickleback ZP proteins, we find that only ZP3 is glycosylated. Incubation of stickleback egg coats with a *N*-glycanase resulted in loss of ZP3 carbohydrate staining by SDS-PAGE, and a reduction in its apparent molecular weight of ∼4 kDa (Figure 2). This relatively large mass is unusual for an *N*-linked glycan in vertebrates, and may be indicative of a complex, tetra-antennary carbohydrate that could present a recognition surface for sperm (7, 104–106). There is only one potential *N*-linked glycosylation site in stickleback ZP3, ^160^NVS^162^ (Figure 4, modeled in yellow), which correspondingly is the same site found to be *N*-glycosylated in rainbow trout ZP3 (46, 97). In an alignment between rainbow trout and mammalian ZP3 proteins, Darie et al. (97) found that this particular *N*-linked glycosylation site is highly conserved to mammals. While it is interesting to note that ZP1 appears to have lost glycosylation entirely in stickleback, this is consistent with what has been described in other fish (39, 46, 97).

Disulfide bonds play important roles in protein folding and structural stability, particularly for secreted proteins, and are a defining characteristic of ZP module-containing proteins with their 8, 10, or 12 conserved, disulfide bonded cysteines (12, 26, 107). Stickleback egg coats are remarkably insoluble relative to other characterized egg coats, a biochemical feature that seems true of fish egg coats in general (9, 47, 108). For instance, we have found that stickleback egg coats remain intact in the presence of 7 M urea, but dissolve better with the addition of a reducing agent, suggesting that disulfide bonds could contribute to their significant structural stability. Notably, the ZP module of ZP1-like proteins from fish contains two extra cysteine residues in a linker between the ZP-N and ZP-C domains (see Figure 3; (97, 109)). This interdomain linker has been implicated in homo- and heterodimeric assembly of ZP proteins, and it is possible that these additional cysteines play a role in fish egg coat stability (20, 21). To determine the pattern of disulfide bonding in stickleback ZP proteins, egg coats were treated with differing reduction and alkylation conditions prior to performing LC-MS/MS with dynamic exclusion turned off, to allow more quantitative peptide spectral counting. We found a consistent pattern of alkylatable cysteines present in stickleback egg coats, as detected by carbamidomethyl (CAM) modification of these residues by mass spectrometry (Figure 3, Tables 1 and 2). Typically these cysteines would be expected to participate in disulfide bonds, and should not be modifiable without reduction. Free cysteines suggest the potential for disulfide shuffling throughout the stickleback egg coat – in fact, nearly all cysteines in ZP1 and ZP3 were found to be CAM modified at least some of the time (Figure 3, denoted by dashed disulfide bonds). It is not clear whether these labile disulfide bonds are intra- or intermolecular, but free cysteines imply structural flexibility in the disulfide bonding of stickleback egg coats. Potential disulfide shuffling is especially apparent in the ZP-N domains of ZP1 and ZP3, the region of the ZP module known to be involved in ZP protein polymerization (16, 110). There are an odd number of cysteine residues in both stickleback ZP1 and ZP3, consistent with the proposed prevalence of disulfide shuffling.

In teleost fish, ZP genes are known to exhibit both ovarian and hepatic expression (9, 23, 47–49). To determine the site of ZP synthesis in stickleback, primers were designed against *ZP1* and *ZP3* and amplified from both ovary and liver cDNA. ZP primers amplified transcripts from liver cDNA only, suggesting that in stickleback these genes are transcribed in the liver (Figure S3). The secreted protein products then make their way through the bloodstream to the ovary, where they assemble around developing oocytes. Both stickleback ZP1 and ZP3 have lost their canonical transmembrane domain (TMD), in agreement with this altered biosynthesis pattern. Although stickleback ZP proteins lack a TMD, they retain a C-terminal hydrophobic region typical of ZP proteins.

The polymerization of ZP proteins into the higher order structure of the egg coat is best characterized in the mouse, where the egg coat matrix consists of heterodimers of ZP2 and ZP3 that polymerize non-covalently into long fibrils interconnected by cross-links of ZP1 (25, 100, 111). While intramolecular disulfide bonds stabilize the native conformation of secreted ZP proteins, the mouse egg coat matrix also contains intermolecular disulfide bonds in the form of cross-linking ZP1 homodimers (20, 34, 112–114). Both ZP2 and ZP3 are required for egg coat formation, as ZP2 or ZP3 knockout mice fail to produce egg coats (114, 115). ZP1 knockout mice do form an egg coat, but it is loose and not interconnected and females are less fertile than wild-type (115, 116). It is interesting to note that ZP4 – a ZP1 homolog pseudogenized in mouse – can be substituted in place of ZP2 in transgenic mice so that ZP3/ZP4 heterodimers form the egg coat matrix rather than ZP2/ZP3 (33). This agrees with the observation that the structural function of ZP2 in mammals is performed by ZP1-like subunits in fish, which lack ZP2 (20, 26, 49, 117). It is also consistent with our finding that ZP1 and ZP3 constitute the stickleback egg coat matrix. ZP-N domains within ZP proteins are thought to facilitate egg coat polymerization, with cross-linking between filaments mediated by ZP1 (7, 16, 28).

As alluded to above, there are interesting changes in stickleback ZP protein architecture relative to what is known about other ZP proteins. Classical ZP protein architecture consists of a N-terminal signal sequence (SS) that marks them as secreted proteins; potential sequence upstream of the ZP module containing additional ZP-N domain repeats, or a P/Q rich-region and trefoil domain in ZP1-like proteins; the ZP module, with its paired ZP-N and ZP-C domains; a consensus furin cleavage site (CFCS) that allows cleavage of the C-terminal region; and a hydrophobic region or TMD (26–28). Changes to stickleback ZP protein architecture are highlighted in approximate order from N- to C-terminus (see Table 4 for summary). First, ZP1 proteins typically have a single N-terminal ZP-N domain repeat upstream of the ZP module, which stickleback ZP1 has lost (18). Stickleback ZP1 has also lost its fourth canonical cysteine in the ZP-N domain of its ZP module (Figure 3). Stickleback ZP1 has two additional cysteine residues, C_4_ and C_5_, in a linker between the ZP-N and ZP-C domains of its ZP module that are specific to fish (Figure 3, boxed in black; (26, 97, 109)). Stickleback ZP3 also has an additional cysteine residue, C_9_, in its ZP-C domain (Figure 3, boxed in black). Both stickleback ZP proteins have lost their TMDs, likely as a consequence of their hepatic expression (26, 39, 118). Stickleback ZP1 appears to have lost all glycosylation, while stickleback ZP3 contains a single *N*-linked glycan in the linker between its ZP-N and ZP-C domains at a site well-conserved from fish to mammals (Figure 4, modeled in yellow; (39, 46, 97)). Finally, disulfide shuffling is prevalent in both stickleback ZP proteins, particularly within the ZP-N domains of their ZP modules, and particularly for ZP1 (see Figure 3). All cysteines in the ZP-N domain of stickleback ZP1 were modifiable with iodoacetamide in the absence of reducing agent, whereas only C_1_ and C_4_ of ZP3 were – C_2_ and C_3_ formed a stable disulfide bond.

**Table 4:**
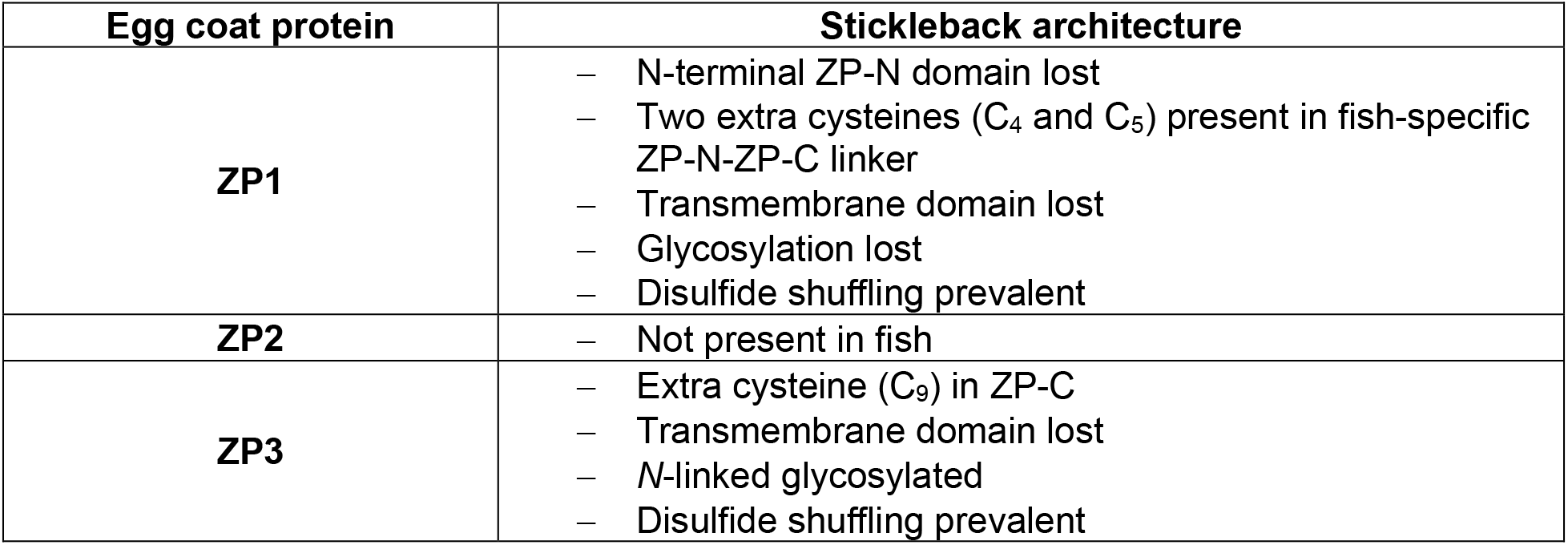
Summary of changes to stickleback ZP protein architecture. Summary of changes to stickleback ZP protein architecture relative to mammalian ZP proteins.

The role of the ZP-N domain in protein polymerization is not limited to reproductive proteins, and is conserved throughout eukaryotes (16, 17, 20, 21, 119). ZP-N/ZP-N interactions between ZP3 and ZP1/2/4 (depending on which ZP proteins are present) are thought to assemble into the structure of the egg coat, so it is notable that stickleback ZP1 has lost one of its two ZP-N domains with the loss of its canonical N-terminal ZP-N repeat. Similarly, ZP2, with its numerous N-terminal ZP-N repeats, is not found in fish (20, 26, 49, 117). Stickleback egg coats may compensate for the loss of these ZP-N polymerization domains with intermolecular, covalent disulfide cross-links arising from disulfide shuffling, which would be a departure from what has been characterized in other animals. The absence of a TMD in stickleback ZP proteins, and often in fish ZP proteins more generally, suggests that the topology of ZP proteins during egg coat assembly may be different in fish relative to mammals as well (46).

The evolution of stickleback *ZP3* under positive Darwinian selection also has interesting implications for stickleback egg coat architecture. In general, rapid evolution is a hallmark of reproductive proteins (51, 120). Numerous evolutionary forces have been attributed to the rapid evolution of reproductive proteins, including sperm competition, sexual conflict (at the cellular level, cryptic female choice), reinforcement, and pathogen resistance (51, 54, 120–122). Using a maximum likelihood method to assess ZP protein evolution across teleost fish, we find that *ZP3* has been subjected to positive Darwinian selection along the lineage while *ZP1* has not (Table 3). Rapid evolution in *ZP3* has also been found in mammals (54, 98, 99). Stickleback ZP3 has nine rapidly evolving residues, six that fall within its ZP-N domain and three that fall within its ZP-C domain (Figure 4, denoted with red spheres; see also Table 3); notably, as few as ten amino acid changes in a sea urchin reproductive protein can lead to gametic incompatibility (123). Although it is interesting that stickleback *ZP1* has not experienced positive selection in teleosts, studies of mammalian *ZP1* similarly find no evidence of positive selection. ZP1 is thought to play a cross-linking role in mammalian egg coats, and stickleback ZP1 may be serving a similar structural function with its parallel evolutionary trajectory. We see many changes in stickleback ZP1 relative to other characterized ZP1 proteins, including the prevalence of disulfide shuffling (even relative to stickleback ZP3), the two extra cysteines that may be involved in homo- or heterodimeric ZP assembly, the loss of its N-terminal ZP-N domain, and the loss of glycosylation. These modifications hint at a conserved structural function, whereas stickleback ZP3 could be playing another role besides contributing to egg coat structure that necessitates evolutionary flexibility. In the mouse, ZP3 has been implicated as a receptor for sperm binding (124). *O*-glycans at S_332_ and S_334_ were identified as sperm ligands, although more recent work has demonstrated that these sites lack glycosylation *in vivo* and are tolerant to mutagenesis without affecting fertility, calling into question the hypothesis of ZP3 as the primary mouse sperm receptor (34, 114, 125, 126). Regardless, amino acids in and around this “sperm-combining site” have been identified as under positive Darwinian selection in a diverse set of mammals (54, 98, 124). That *ZP3* is maintained under selection from stickleback to mammals is intriguing. Although a purely structural role has been suggested for fish ZP proteins given the presence of the micropyle in the egg coat, it is possible that the residues under selection in stickleback ZP3 participate in sperm recognition at the micropyle, particularly given the spatial clustering of the loops containing residues under selection (see Figure 4). These loops of positive selection in ZP3 would therefore be adaptive at the micropyle, but neutral in the remainder of ZP3 molecules forming the rest of the egg coat. The importance of fertilization likely creates a strong selective pressure that could drive rapid evolution, even if this rapid evolution has a functional consequence in only a very small percentage of molecules.

Taken together, our results suggest that the egg coats of stickleback fish are a uniquely protective structure relative to mammalian egg coats. Whereas mammalian sperm secrete acrosomal proteins to bind to the egg coat and create a hole at the point of contact, fish sperm lack an acrosome and enter the egg coat through a specialized channel, the micropyle (7, 39, 42, 59). It is conceivable that the presence of this structure has favored evolutionary events leading to an otherwise impenetrable egg coat: freed from the need to permit sperm access via transient, reversible ZP-N/ZP-N interactions, stickleback egg coats have evolved covalent cross-links arising from disulfide shuffling to stabilize the matrix. Given that fish eggs develop in external environments, such as the bottom of a lake or ocean, subject to high levels of mechanical stress – as well as potential pathogen exposure – a protective structural barrier might be evolutionarily favored (6, 7). Another mechanism for building impenetrable egg coats involves covalent cross-linking of ZP proteins via the N-terminal P/Q-rich region of ZP1, by the action of a transglutaminase enzyme (9, 26, 46, 97, 108). These heterodimeric cross-links would not be reversed under reducing conditions, however, and so are unlikely to represent a significant contribution to egg coat structural stability the way intermolecular disulfide bonds in stickleback are. Correspondingly, only small amounts of these P/Q cross-linked heterodimers are detected by mass spectrometry in unfertilized rainbow trout eggs (46). On the other hand, these transglutaminase cross-links are likely important after fertilization, where they harden the egg coat to further reinforce the matrix and block polyspermy (7, 9, 26, 28, 39, 47).

In summary, there are unique biochemical attributes of fish ZP proteins that likely create a different set of protein-protein interactions for egg coat assembly and fertilization than has been characterized in other animals. The structure of the micropyle may underlie these changes. In teleost eggs, the inner micropylar opening directly adjoins the egg plasma membrane, creating what may be a specialized site for binding fertilizing sperm (42). The recently described zebrafish egg plasma membrane protein Bouncer – which permits cross-species fertilization between medaka and zebrafish, separated by 200 million years of evolution, expressing the medaka version of Bouncer – represents a possible candidate for sperm recognition at the egg plasma membrane (127). Our findings suggest that ZP3 in the egg coat may also contribute to sperm recognition at the micropyle, given its suggested role as a sperm receptor in mammals and its maintenance under positive Darwinian selection from teleost fish to mammals.

## Acknowledgements

We thank Dr. Catherine Peichel (Fred Hutchinson Cancer Research Center, Seattle, Washington, USA and University of Bern, Bern, Switzerland) for her expertise on stickleback biology and assistance with fish collection and Mari Kawaguchi (Sophia University, Tokyo, Japan) for providing *ZP1* and *ZP3* gene sequences for primer design. This work was supported by NIH fellowships T32 HG000035-22 and F31 HD093441 (to E.E.K.), K99 HD090201 (to D.B.W), and NIH grant R01 HD076862 and UW Royalty Research Fund A111769 (to W.J.S.). The content is solely the responsibility of the authors and does not necessarily represent the official views of the National Institutes of Health.

## Data Availability

The mass spectrometry proteomics data have been deposited to the ProteomeXchange Consortium via the PRIDE partner repository with the dataset identifiers PXD017488 (disulfide characterization data) and PXD017489 (egg coat mass spectral characterization data) (128).

## Additional Information

The authors declare that they have no conflicts of interest with the contents of this article.

## Abbreviations

BEB: Bayes empirical Bayes
C/Cys: cysteine
C-terminal: carboxy-terminal
CAM: carbamidomethyl modification
CFCS: consensus furin cleavage site
*d_N_*: number of nonsynonymous substitutions per nonsynonymous codon site
*d_S_*: number of synonymous substitutions per synonymous codon site
IAA: iodoacetamide
N-terminal: amino-terminal
NSAF: normalized spectral abundance factor
ω: *d_N_*/*d_S_*
P/Q: proline/glutamine
S/Ser: serine
SS: signal sequence
TMD: transmembrane domain
ZP: zona pellucida
ZP-C: C-terminal domain of ZP module
ZP-N: N-terminal domain of ZP module

---

**Table S1:**
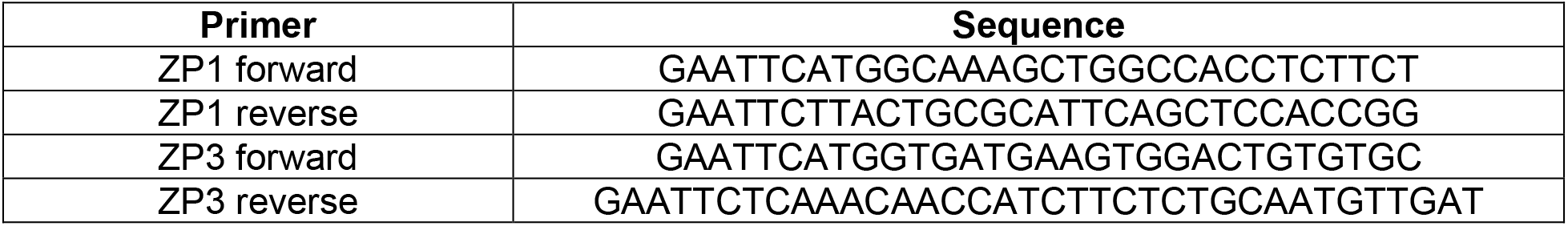
Gene-specific primers used for ZP RT-PCR. Primers were designed to anneal to the 5’ and 3’ UTR of stickleback ZP1 and ZP3, and included an EcoRI overhang (GAATTC).

**Table S2:** Summary of mass spectral data for SDS-PAGE-extracted stickleback egg coat proteins. Table of LC-MS/MS identifications for gel extracted stickleback egg coat proteins (see Figure S2). Sequence matches were sorted by normalized spectral abundance factor (NSAF), with the most abundant Trinity contig match reported (129). Note that for Gel Band 3, the two Trinity contig matches have identical protein sequence.

| Gel band number | Accession ID(s) | Sequence match | Number of peptides | Number of spectra | Percent total coverage | NSAF |
| --- | --- | --- | --- | --- | --- | --- |
| 1 | TR556\|c0_g1_i1_1+ | Vitellogenin | 62 | 296 | 76 | 0.0038521 |
| 2 | TR556\|c0_g1_i1_1+ | Vitellogenin | 89 | 726 | 78 | 0.087787 |
| 3 | TR6893\|c1_g15_i1_3-<br>TR6893\|c1_g17_i1_1- | ZP1 | 38 | 329 | 54 | 0.012559 |
| 4 | TR7203\|c0_g14_i1_3+ | ZP3 | 39 | 244 | 50 | 0.022084 |
| 5 | TR7203\|c0_g14_i1_3+ | ZP3 | 56 | 916 | 54 | 0.035461 |
| 6 | – | Unknown | – | – | – | – |
| 7 | TR7069\|c0_g6_i1_1- | Vitellogenin | 51 | 220 | 44 | 0.023192 |

**Table S3:**
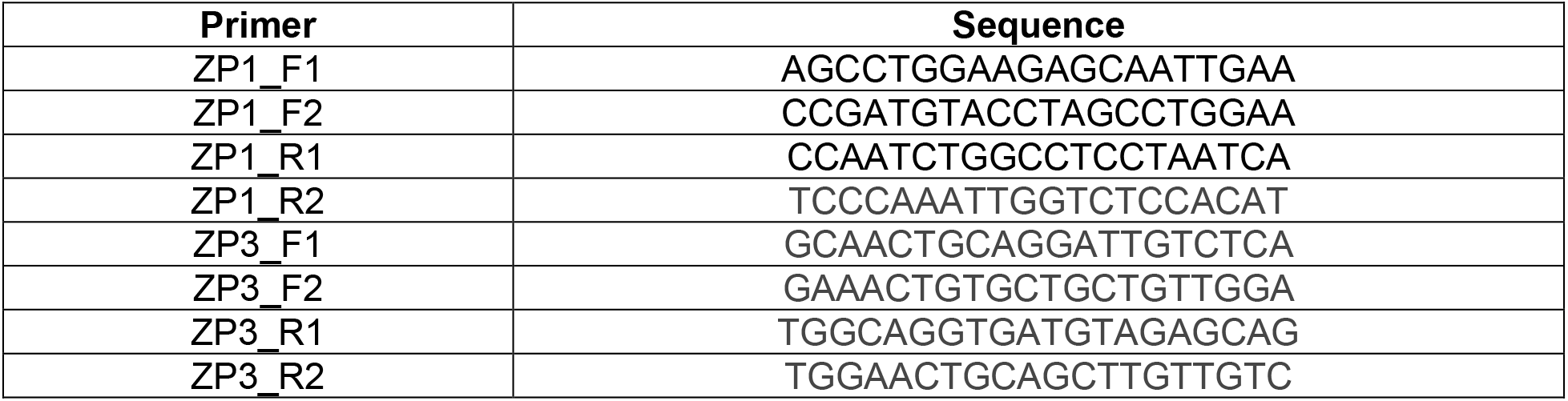
Gene-specific primers used for ZP ovary and liver PCRs. Primers were designed against the most abundant Trinity contig by LC-MS/MS NSAF.

**Table S4:** Expected product sizes for ZP primer pairs. Expected product size for each ZP primer pair (see Figure S3).

| Lane | Forward primer | Reverse primer | Expected product size (bp) |
| --- | --- | --- | --- |
| 1 | ZP1_F1 | ZP1_R1 | 579 |
| 2 | ZP1_F1 | ZP1_R2 | 725 |
| 3 | ZP1_F2 | ZP1_R1 | 590 |
| 4 | ZP1_F2 | ZP1_R2 | 716 |
| 5 | ZP3_F1 | ZP3_R1 | 669 |
| 6 | ZP3_F1 | ZP3_R2 | 612 |
| 7 | ZP3_F2 | ZP3_R1 | 724 |
| 8 | ZP3_F2 | ZP3_R2 | 647 |

**Figure S1:**
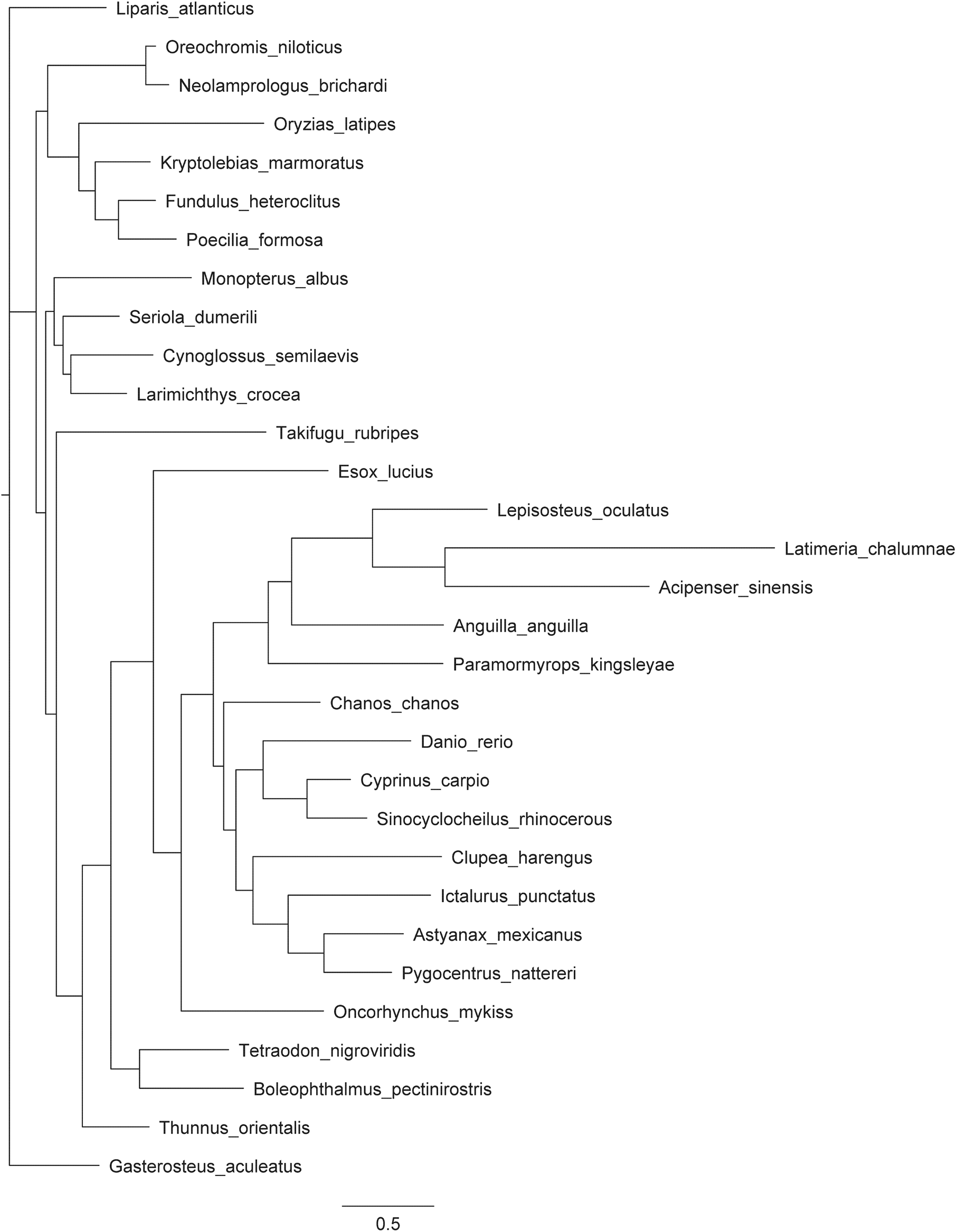
ZP evolution within teleosts. Maximum likelihood evolutionary tree of concatenated *ZP1* and *ZP3* from representative teleost species used for molecular evolutionary analyses.

**Figure S2:**
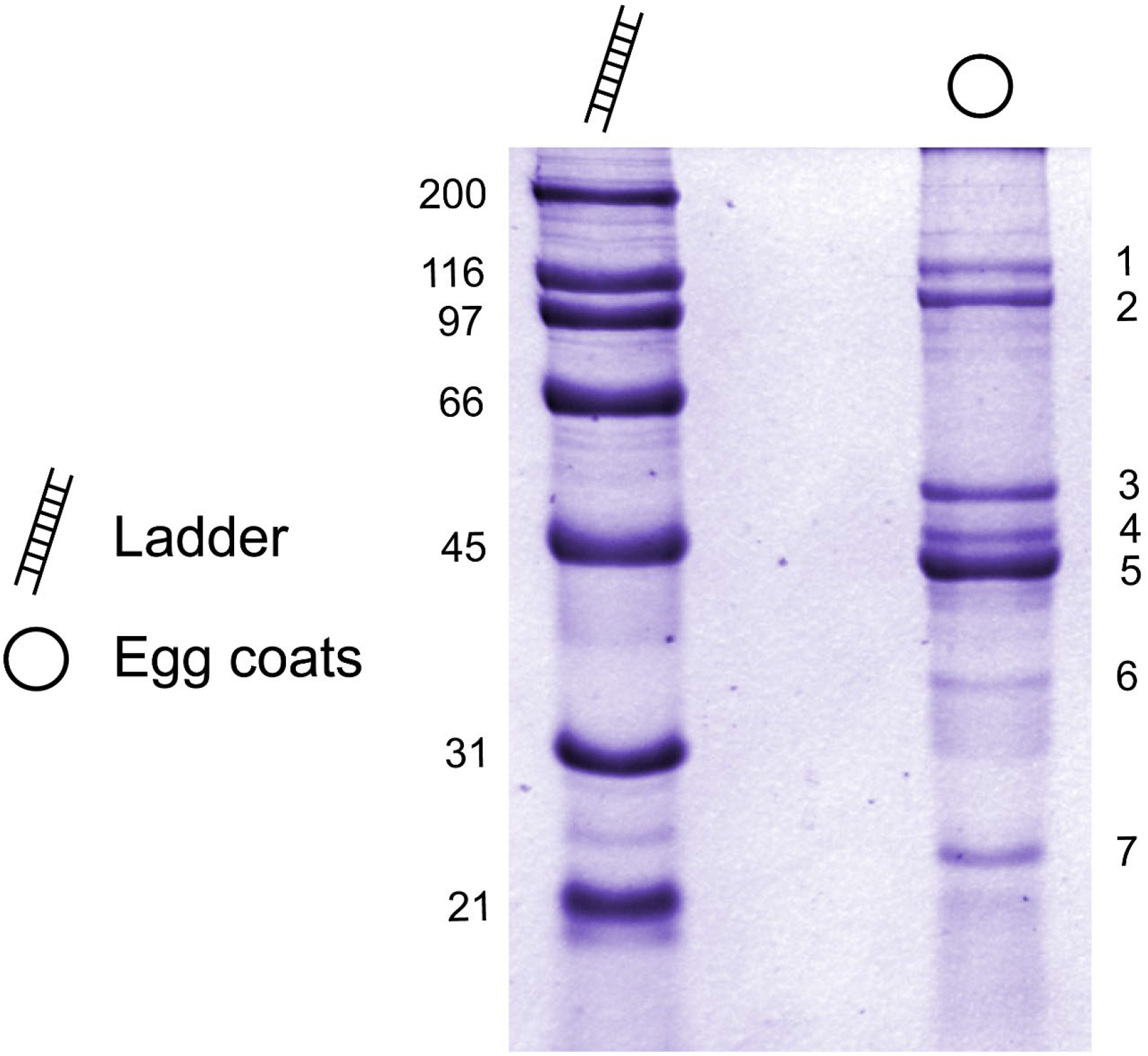
SDS-PAGE of stickleback egg coats for gel extraction-mass spectrometry. Stickleback egg coats were separated with SDS-PAGE, and individual bands (indicated by number, see Table S2) were gel extracted and analyzed by LC-MS/MS. ZP1 and ZP3 were found to be the two major components of stickleback egg coats, with the remaining bands representing vitellogenin contamination likely carried over from egg coat isolation.

**Figure S3:**
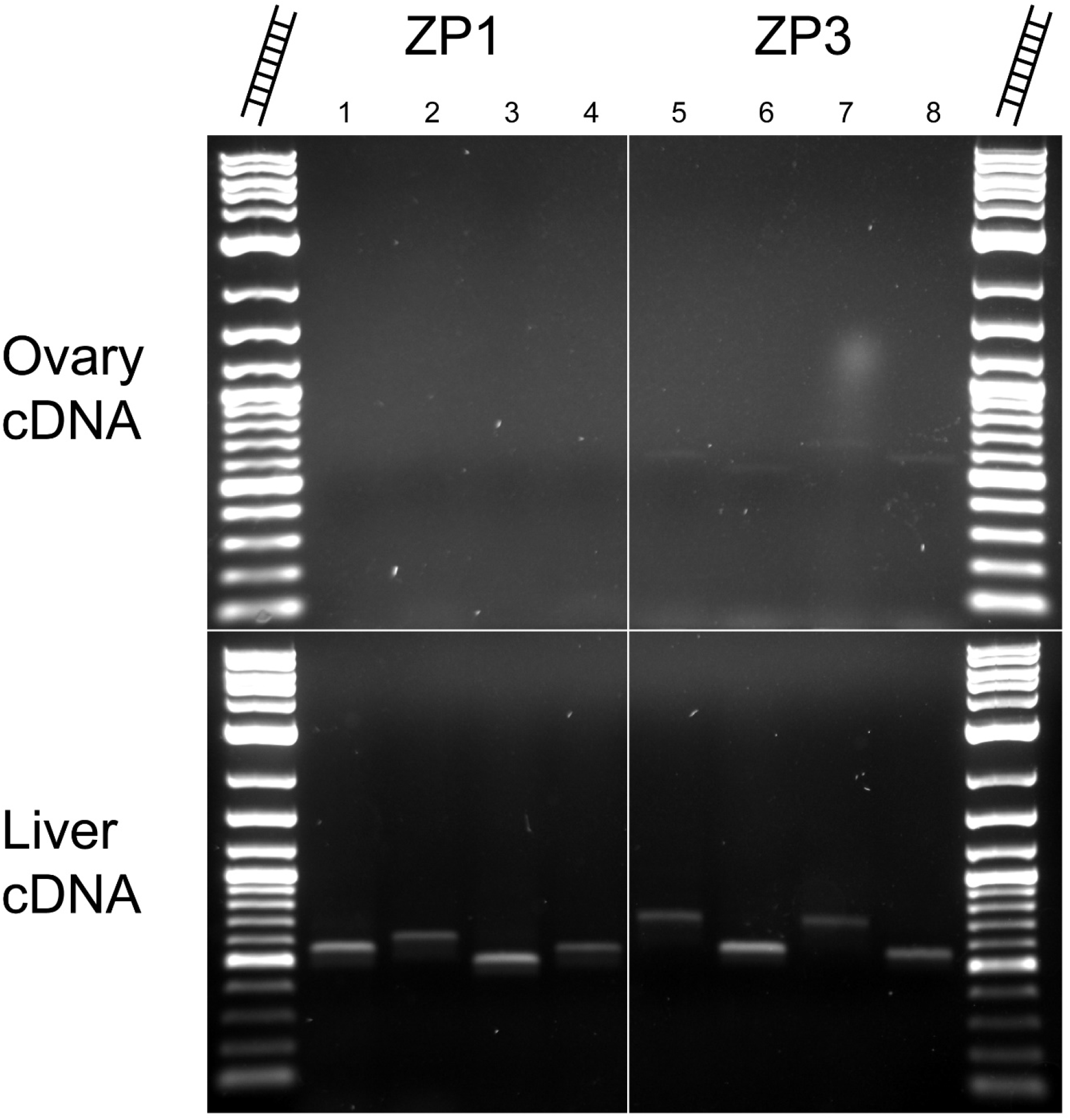
Amplification of ZP1 and ZP3 from stickleback liver cDNA. To determine the site(s) of ZP expression in stickleback, primers were designed against ZP1 and ZP3 (primer sequences in Table S3, expected product size by lane in Table S4) and amplified from stickleback liver and ovary cDNA. Successful amplification occurred only from liver cDNA, suggesting that in stickleback, ZP genes are transcribed in the liver.

